# Temporally resolved glutamate and GABA responses measured in human medial frontal cortex during working memory encoding and recall

**DOI:** 10.64898/2026.02.13.705729

**Authors:** Daniel Cocking, Ryan Elson, Katherine Dyke, Mohammad Zia Ul Haq Katshu, Adam Berrington, Claudia Danielmeier

**Affiliations:** Sir Peter Mansfield Imaging Centre, School of Physics and Astronomy, University of Nottingham, Nottingham, United Kingdom; School of Psychology, University of Nottingham, Nottingham, United Kingdom; Institute of Mental Health, School of Medicine, University of Nottingham, Nottingham, United Kingdom; Nottinghamshire Healthcare NHS Foundation Trust, United Kingdom

**Keywords:** functional magnetic resonance spectroscopy (fMRS), glutamate dynamics, GABA, working memory, medial frontal cortex, event-related design, 7T, cognitive neurochemistry

## Abstract

Resolving rapid neurochemical changes during cognition is key to linking neurophysiology with behaviour. We developed a mixed block/event-related design of a working memory (WM) task for functional magnetic resonance spectroscopy (fMRS) at 7T to resolve glutamate and GABA fluctuations during encoding and recall phases of WM. Participants performed the WM task interleaved with rest and control conditions while semi-LASER spectra were acquired from the medial frontal cortex. Group-level phase-specific analyses revealed significant glutamate, but not GABA, increases during encoding and recall relative to control. However, larger individual increases in GABA during encoding predicted higher WM accuracy. Glutamate exhibited distinct, temporally resolved peaks for encoding (∼1.1s) and recall (∼1.4s). These time courses extend beyond previously reported post-stimulus windows and align closely with behavioural response latencies. Our findings provide temporally high-resolved characterisations of glutamate responses during distinct phases of WM and highlight differential roles of glutamate and GABA in WM processes.

## 1. Introduction

The cognitive processes underlying working memory (WM) emerge from rapid fluctuations in excitatory and inhibitory neural activity, yet these dynamics are challenging to quantify *in vivo*. Probing these dynamic processes in humans offers a critical link between neural activity and behaviour. Using proton (^1^H) magnetic resonance spectroscopy (MRS), resting concentrations of the major excitatory and inhibitory neurotransmitters, glutamate (Glu) and gamma-aminobutyric acid (GABA), have been associated with varied measures of cognitive function^1–5^. Time-resolved measurements during functional tasks (fMRS) have revealed transient changes in Glu and GABA concentrations to cognitive demands ^6–8^, allowing more direct associations with cognition. Such studies typically rely on blocked-designs, averaging measurements over long durations (>15 s) of a task paradigm ^9–17^. This approach limits the ability to assign neurochemical changes to specific cognitive processes within trials, such as the different phases within working memory (WM) paradigms. Although tracking changes to individual events on a sub-second scale with time-locking to stimulus and event-related designs is feasible ^6,18,19^, short timescale temporal dynamics of Glu and GABA, which may reflect synaptic activity ^20^, remain elusive.

Canonical ‘time courses’ of Glu and GABA concentration are currently lacking. Glu has been proposed to peak in concentration around ∼0.5 s after stimulus onset ^17^ and return to baseline after three to four seconds, ^17,19^. The lack of an established response complicates study design in determining the optimal time to acquire spectroscopy data post-stimulus. Compared to the postulated peak at 0.5 s ^17^, some prior work may have sampled too early, ^6,21–23^ or too late ^24^. Small concentration changes make measuring time courses of neurochemicals difficult, thus fMRS studies often measure changes over long durations. Cognitively demanding tasks such as WM paradigms, however, offer a possibility to elicit strong responses, and hence detectable signal changes, over timescales on the order of single events.

Working memory is a crucial cognitive ability allowing us to attend to and manipulate information held in short-term memory ^25^. It comprises of three core phases: encoding, maintenance and retrieval. A commonly used WM paradigm is the n-back task ^7,8^ which conflates encoding and recall, obscuring which neurochemical shifts occur during different phases of WM. Therefore, to characterise the temporal dynamics of WM-related neurochemical responses with fMRS in different phases, a task paradigm separating encoding, maintenance, and retrieval is required.

In the current work, we developed a unique event-related fMRS acquisition at 7T, sensitive to the temporal dynamics of Glu by sampling at 16 time points over a 2.5 s window post-stimulus. Our experimental design allowed separate estimation of Glu concentration time courses associated with encoding and recall phases of an associative WM task. Participants encoded the colour, shape and location of four abstract shapes, then recalled the colour from a shape or location cue. Normalising to control trials allowed us to isolate WM-specific processes over and above motor and visual responses within an fMRS voxel situated in the pre-SMA region of medial frontal cortex (MFC), guided by a separate fMRI localiser study (see Supplementary Materials). The MFC is involved in many of the cognitive functions required for associative WM, such as executive function and attention ^26–30^. Different subpopulations of neurons in the pre-SMA region have also been associated with different stages of WM ^31^.

We found evidence for a transient Glu response in the MFC of healthy human participants at 7T, displaying peak increases in concentration around 1.1s and 1.4s, following encoding and recall, respectively. At an individual level, Glu change was not related to task performance, whereas changes in GABA concentrations compared to control trials were positively correlated with WM performance, particularly during encoding, suggesting that inhibitory processes, potentially to inhibit task-irrelevant information, are important during demanding WM tasks.

## 2. Results

### 2.1. Time courses of Glu reveal distinct peaks during encoding and recall

We estimated Glu time courses from our MRS voxel during WM encoding and recall and in control trials. Across the trials, data were acquired at 16 sampling points between stimulus onset (0 s) and stimulus offset (2.5 s) in 0.17 s intervals. To improve neurochemical estimation for Glu, spectra were binned into eight timepoints prior to spectral fitting (see Fig. 1). Within each run, concentrations for each time bin were converted to a percentage change from the run’s initial rest period (the “baseline”, comprising eight spectra with Cramer-Rao Lower Bounds [CRLBS] = 9.1% [SE = 0.8%]), and then averaged across runs.

**Figure 1.**
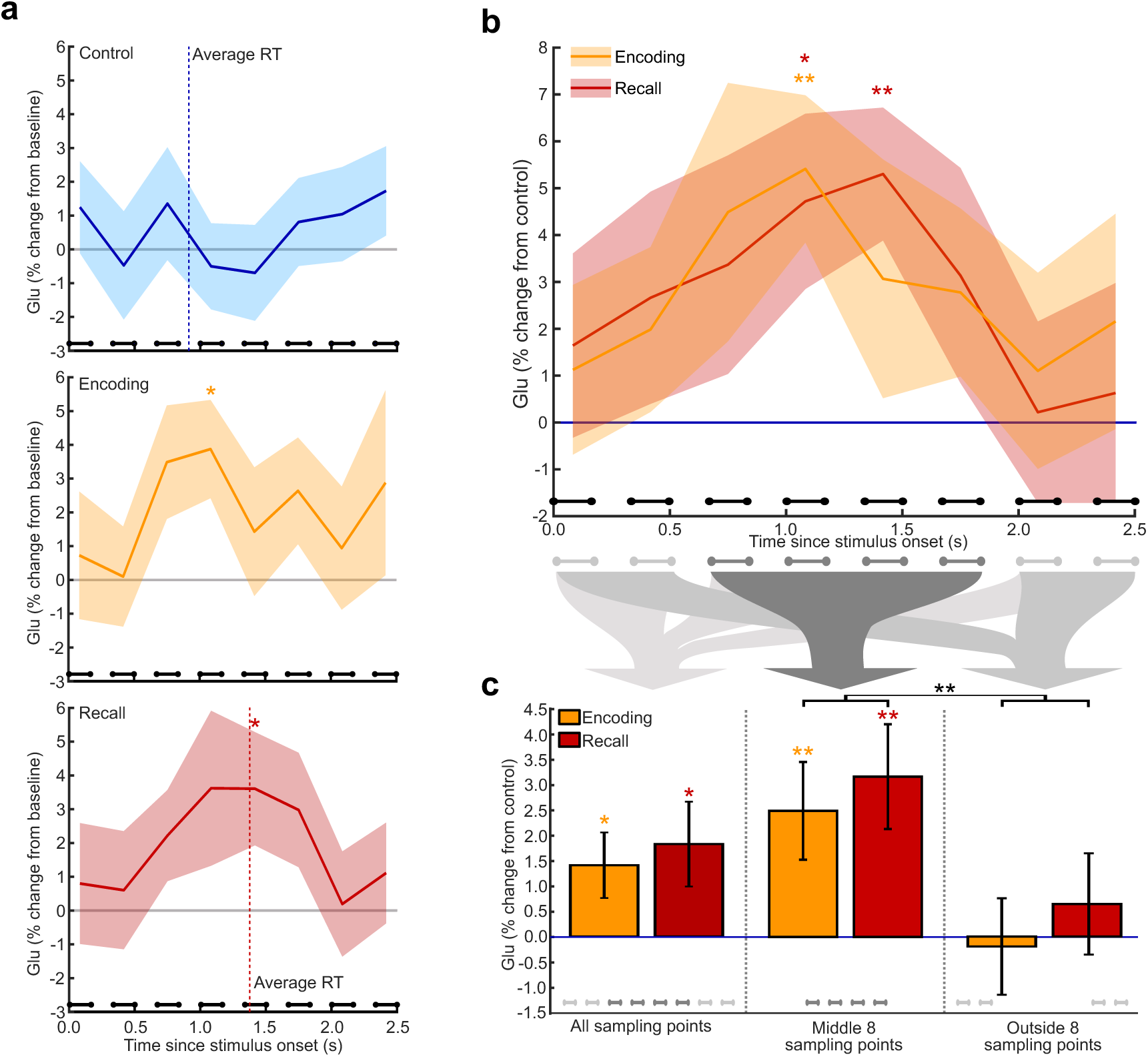
Time courses of Glu concentration changes for Control, Encoding and Recall phases. *Note:* (**a**) Glu time course measured during control trials (blue, top), encoding (orange, middle) and recall (red, bottom) phase as percentage change from the baseline. Vertical dotted lines represent the average reaction times for control (blue, 0.91 s) and working memory trials (red, 1.38 s). Linked dots at the bottom of each graph represent the pair of sampling points in each time bin. (**b**) Glu response for encoding (orange) and recall (red) normalised to control Glu (blue) for each time bin separately (% change). (**c**) average % increases in Glu compared to control trials during encoding (orange) and recall (red) across all 16 sampling points (left), the ‘peak’ window including the middle eight sampling points (middle) and the outside window including the first four and last four sampling points (right). All error bars and shaded regions show the standard error across participants. * Significant uncorrected, ** significant Bonferroni-corrected.

During encoding, Glu was significantly increased compared to baseline, prior to correction, at 1.08 s post stimulus onset (4^th^ time bin [1.00-1.17 s]; mean [SE] = 3.87% [1.45%], t(18) = 2.67, *p* = .016, 95% CI = [0.82, 6.93], *d* = 0.61, see Fig.1a).

During recall, Glu levels were significantly increased, prior to correction, at 1.42 s post probe onset (5^th^ time bin [1.33-1.50 s]; mean [SE] = 3.61% [1.68%], t(18) = 2.15, *p* = .045, 95% CI = [0.09, 7.13], *d* = 0.49). No other timepoint was significant for either WM phase or the control task. Bonferroni-corrected *α* = .002.

During encoding, Glu was significantly increased compared to control at 1.08 s (4^th^ time bin; mean [SE] = 5.41% [1.57%], t(18) = 3.44, *p* = .003, 95% CI = [2.11, 8.71], *d* = 0.79), measured by converting raw concentrations during encoding (and recall) to percent changes relative to control, demonstrating a Glu increase specific to WM-related processes, independent from visual or motor demands in control trials (Fig. 1b). This normalisation was performed separately for each time point, with changes averaged over runs.

During recall, Glu was significantly increased compared to control at 1.42 s (5^th^ time bin; mean [SE] = 5.30% [1.42%], t(18) = 3.74, *p* = .002, 95% CI = [2.32, 8.28], *d* = 0.86) and, prior to correction, was also increased at 1.08 s (4^th^ time bin; mean [SE] = 4.72% [1.87%], *t*(18) = 2.52, *p* = .021, 95% CI = [0.79, 8.65], *d* = 0.58). Bonferroni-corrected *α* = .003.

The mean (SE) Glu Cramér-Rao Lower Bound (CRLB) values across each time point for control, encoding and recall WM phases were 12.25% (1.24%), 12.05% (1.16%) and 12.01 (1.11%) respectively. CRLBs did not significantly differ by time point or phase indicating no difference in data quality.

### 2.2. Glu, but not GABA, increases during encoding and recall

The novel time course reconstructions allowed us to interrogate the Glu response over time. We employed subsequent analyses to compare the overall Glu and GABA concentration changes averaged over longer post-stimulus time windows (two windows ∼1.25 s, one 2.5 s). To isolate WM-specific changes in neurochemical concentrations, results below are reported as the percentage change from control trials.

We aimed to test three questions. 1) Did the concentration changes during encoding and recall differ from control trials? 2) Did the concentration changes differ between encoding and recall? 3) Were the concentration changes significantly correlated across encoding and recall?

On average, Glu increased by 1.40% (SE: 0.65%) during encoding and 1.82% (SE: 0.84%) during recall relative to control (Fig. 1c). This was shown by initially averaging spectra for all 16 sampling points prior to fitting (32 spectra per fit per run) These increases were significant prior to Bonferroni correction (*α* = .025, encoding: *t*(18) = 2.17, *p* = .044, 95%CI = [0.05, 2.76], *d* = 0.50, recall: *t*(18) = 2.18, *p* = .043, 95%CI = [0.06, 3.58], *d* = 0.50). Concentration changes during encoding and recall did not significantly differ (*t*(18) = 0.55, *p* = .591, 95%CI = [-1.18, 2.02], *d* = 0.13) and were significantly correlated (*r* = 0.50, *p* = .030). The mean (SE) Glu CRLB values did not significantly differ between control, encoding and recall WM phases: 5.72% (0.44%), 5.66% (0.41%) and 5.65 (0.42%)

#### 2.2.1. Windowing (0.67-1.83 s) increases the Glu response

Given the inverted U-shaped rise and fall in the time course of Glu, we employed a windowed averaging approach centred around the peaks (∼1.08 and ∼1.42 s) of the Glu response, assessing Glu in a critical window without the influence of earlier and later time points where the concentration appears to be at baseline.

We separated the ‘peak’ (middle) eight sampling points (0.67-1.83 s) focusing on the peak of Glu responses and the ‘outside’ eight sampling points (the first four and last four sampling points). For each analysis, spectra were averaged across the eight sampling points prior to fitting (16 spectra per fit per run; Fig. 1c). ‘Peak’ and ‘outside’ Glu concentrations during encoding and recall were separately normalised to the corresponding windows of the control task (Fig. 1c).

A 2x2 within-subjects ANOVA showed a main effect of time window (peak/outside: *F*(1, 18) = 6.41, *p* = .021, 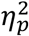 = 0.26), with significantly higher Glu changes in the peak window (mean [SE] = 2.83% [0.77%]) than in the outside time windows (mean [SE] = 0.22% [0.85%]). The effect of WM phase (encoding/recall) was not significant (*F*(1, 18) = 0.92, *p* = .351, 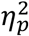 = 0.05) nor was the interaction (*F*(1, 18) = 0.01, *p* = .922, 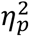 = 0.00). This suggests WM-related Glu changes are restricted to approximately between 0.67-1.83 s post stimulus onset.

In this ‘peak’ time window (0.67-1.83 s), we observed significant Glu increases during encoding (mean [SE] = 2.49% [0.97%], *t*(18) = 2.58, *p* = .019, 95%CI = [0.46, 4.52], *d* = 0.59) and recall (mean [SE] = 3.17% [1.03%], *t*(18) = 3.06, *p* = .007, 95%CI = [1.00, 5.34], *d* = 0.70) compared to the control condition. Bonferroni-corrected *α* = .025. Interestingly, unlike the full window analysis, Glu concentration changes were not significantly correlated across encoding and recall (*r* = 0.17, *p* =.483), suggesting more independent Glu responses related to the different WM phases when the influence of baseline Glu levels is reduced. CLRBs did not significantly differ between control, encoding and recall WM phases using the ‘peak’ sampling points: 6.97% (0.54%), 6.98% (0.57%) and 6.97% (0.51%) respectively.

As the results highlighted the benefits of cropping to the peak time window for Glu, all subsequent analyses, including those involving GABA are also restricted to this middle window. Results incorporating all 16 sampling points can be seen in the Supplementary Materials (e.g., see Supplementary Fig.1).

#### 2.2.2. Group-level GABA concentrations are not modulated during WM phases

GABA concentrations did not significantly differ from control trials during encoding (mean [SE] = -1.11% [3.33%], *t*(17) = -0.33, *p* = .743, 95%CI = [-8.15, 5.92], *d* = 0.08) or recall (mean [SE] = 1.04% [2.41%], *t*(17) = 0.43, *p* = .672, 95%CI = [-4.02, 6.12], *d* = 0.10). Bonferroni-corrected *α* = .025. The concentration changes did not significantly differ between encoding and recall (*t*(17) = 0.66, *p* = .517, 95%CI = [-4.70, 9.00], *d* = 0.16) and were highly correlated (*r* = 0.63, *p* = .004). GABA CLRBs did not significantly differ between control, encoding and recall WM phases: 21.48% (1.40%), 23.01% (1.60%) and 22.52% (1.29%), respectively. The GABA CRLB values are larger than that of Glu analysis, and therefore, to maintain reliability of measurements, spectra were not split into smaller bin sizes.

### 2.3. Changes in GABA, but not Glu, predict WM performance

We investigated the relationship between task performance (percent accuracy and RT) and the change in Glu and GABA levels during encoding and recall (averaged around the Glu peak as described above) relative to control trials.

Correlations can be seen in Fig. 2 (for completeness, correlations using all sampling points can be seen in the Supplementary Materials and Supplementary Fig. 2).

**Figure 2.**
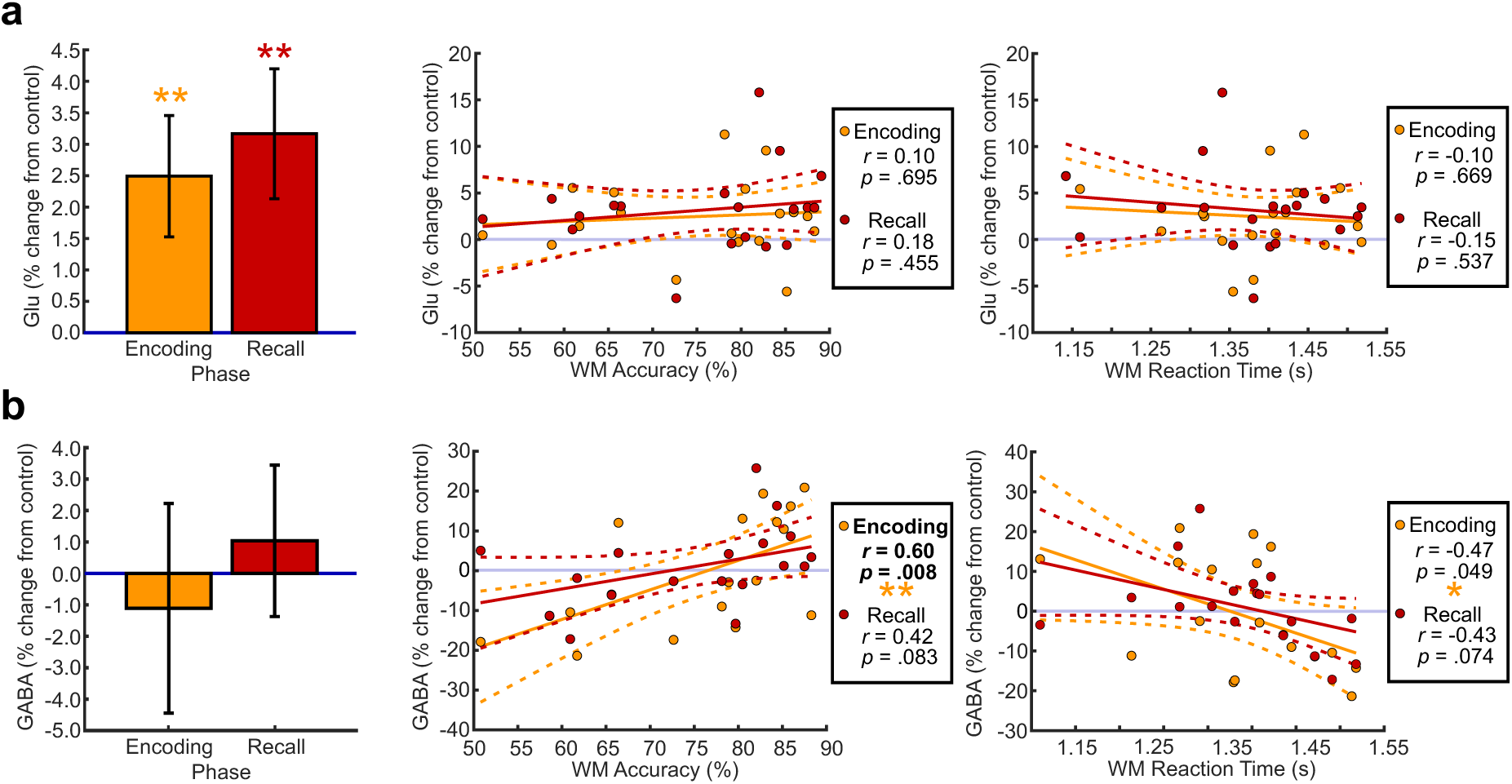
Correlations between the Glu (**a**) and GABA (**b**) changes from control with working memory performance. *Note:* Percent signal change from control trials during encoding (orange) and recall (red), showing changes in Glu (**a**) and GABA (**b**), showing the group-level changes (left) and the correlations with working memory accuracy (middle) and working memory reaction time (right). Plots show data using the middle eight sampling points. * Significant prior to correction, ** significant after correction.

Interestingly, increases in GABA during WM encoding predicted better task performance (Fig. 2b), with participants with more GABA increases showing higher accuracy (*r* = 0.60, *p* = .008; 1 outlier removed with Cook’s distance >1 in all correlations) and faster reaction times, significant prior to correction (*r* = -0.47, *p* = .049). Trends in the same direction were present for GABA during recall (accuracy: *r* = 0.42, *p* = .083, RT: *r* = -0.43, *p* = .074). Bonferroni-corrected *α* = .025. Thus, while group-level GABA concentrations were not significantly increased during WM phases compared to control, individual-level GABA dynamics predicted task performance. In contrast, changes in Glu during encoding or recall were not significantly correlated with accuracy or RT on the WM trials (*all p* >= .455, see Fig. 2a).

### 2.4. Trend towards diminishing Glu increase across runs

To assess the effect of run on neurochemical responses (see Fig. 3), we averaged the percent changes in glutamate and GABA across encoding and recall. See supplementary materials (Supplementary Fig. 3) for results using all 16 sampling points.

**Figure 3.**
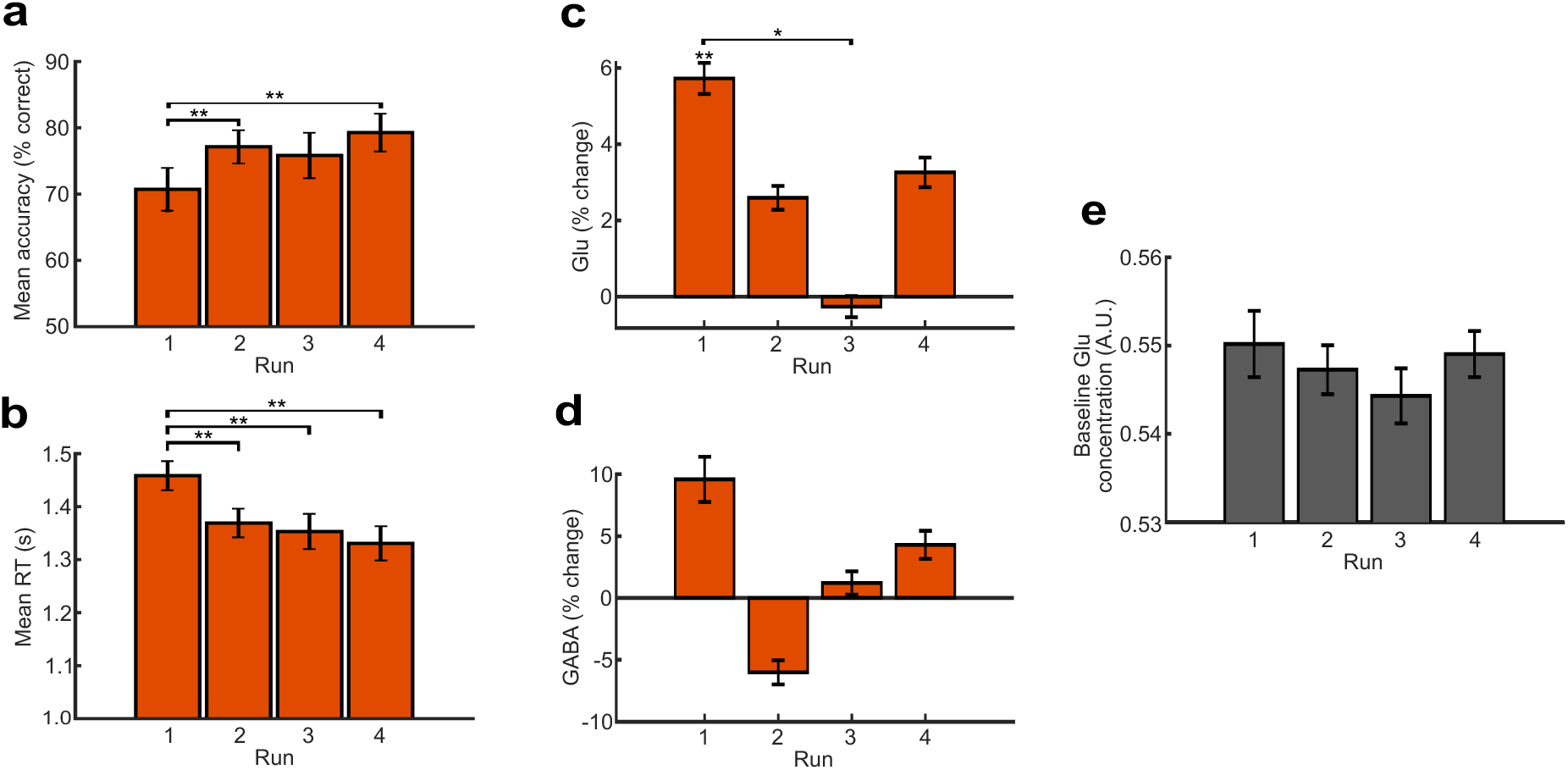
Behavioural data, Glu and GABA across runs. *Note*: Data across runs. (**a**) Mean accuracy and (**b**) reaction time during the working memory trials. Mean percent Glu (**c**) and GABA (**d**) change within memory trials (averaged across encoding and recall) from control trials using the middle eight sampling points. (**e**) Raw Glu concentration at baseline. Error bars show +/- 1 SE across participants. * Significant prior to correction, ** significant after correction.

For Glu, mean percent signal changes (SE) per run were: Run 1: 5.73% (1.78%), Run 2: 2.59% (1.36%), Run 3: -0.27% (1.21%), Run 4: 3.26% (1.70%). The within-subjects ANOVA showed a trending effect of run (*F*(3, 56) = 2.60, *p* = .062, 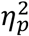 = 0.13, see Fig. 3c). The decrease in Glu change from Run 1 to Run 3 was significant prior to correction (*α* = .008, *t*(18) = -2.57, *p* = .019, 95% CI = [-10.88, - 1.10], *d* = 0.59). No other differences between runs were significant (all *p* >= .102). The assumption of sphericity was met.

Compared to control trials, Glu showed a significant increase for Run 1 only (*t*(18) = 3.23, *p* = .005, 95% CI = [2.00, 9.46], *d* = 0.74). The increases in Run 2 (*t*(18) = 1.90, *p* = .073, 95% CI = [-0.27, 5.45], *d* = 0.44) and Run 4 (*t*(18) = 1.92, *p* = .071, 95% CI = [-0.31, 6.83], *d* = 0.44) only trended towards significance prior to correction (*α* = .013).

For additional context, there was no significant effect of run for the raw Glu concentrations obtained during the initial rest spectra at the start of each run (eight spectra per fit, *F*(3, 54) = 0.07, *p* = .975, 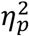 = 0.00, see Fig. 3e) showing that Glu decreases over time were related to participants performing the task but returned to similar concentrations in between runs.

For GABA (Fig. 3d), the effect of run was not significant (*F*(2.01, 36.26) = 1.58, *p* = .220, 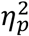 = 0.08, Greenhouse-Geisser correction applied) and concentrations during WM blocks did not differ from control (all *p* >= .175). The mean percent changes were: Run 1: 9.58% (7.96%), Run 2: -6.00% (4.25%), Run 3: 1.21% (4.10%), Run 4: 4.29% (4.93%).

### 2.5. WM performance improves across runs

The mean (SE) accuracy rate across all participants and runs was 75.74% (2.66%) and the mean reaction time (RT) was 1.38 s (0.02 s).

For memory trials, we explored the main effect of run (1-4) to check if performance decreases in line with the trend seen for Glu (Fig. 3c). There were main effects for both accuracy (*F*(3, 54) = 4.80, *p* = .005, 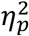 = 0.21) and reaction time (*F*(3, 54) = 7.11, *p* < .001, 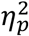 = 0.28).

Mean accuracies (SE) per run were: Run 1: 70.72% (3.23%), Run 2: 77.14% (2.51%), Run 3: 75.82% (3.42%), Run 4: 79.28% (2.85%). Accuracy was significantly higher in Run 2 (*t*(18) = 3.17, *p* = .005, 95% CI = [2.17, 10.66], *d* = 0.73) and Run 4 (*t*(18) = 3.02, *p* = .007, 95% CI = [2.59, 14.51], *d* = 0.69) compared to Run 1. The difference between Runs 1 and 3 trended towards significance prior to correction (*t*(18) = 2.05, *p* = .055, 95% CI = [-10.32, 0.12], *d* = 0.47). No other comparisons were significant. Bonferroni-corrected *α* = .008. See Fig. 3a.

Mean RTs per run were: Run 1: 1.46 s (0.03 s), Run 2: 1.37 s (0.03 s), Run 3: 1.35 s (0.03 s), Run 4: 1.33 s (0.03 s). Reaction times were significantly faster in Run 2 (*t*(18) = -3.95, *p* < .001, 95% CI = [-0.14, -0.04], *d* = 0.91), Run 3 (*t*(18) = -3.05, *p* = .007, 95% CI = [-0.18, -0.03], *d* = 0.70), and Run 4 (*t*(18) = -3.37, *p* = .003, 95% CI = [-0.21, -0.05], *d* = 0.77) compared to Run 1. No other comparisons were significant. Bonferroni-corrected *α* = .008. See Fig. 3b.

Behavioural data in control trials was at ceiling and therefore not included in any analyses above. Mean (SE) accuracy was 98.48% (1.02%) and reaction time was 0.91 s (0.04 s).

## 3. Discussion

Functional MRS studies offer new insights into the dynamics of neural processes that occur during cognitive tasks. Here, we developed an event-related fMRS approach to characterise dynamic neurochemical changes in the posterior MFC during an associative WM task. Glutamate concentration measured post-stimulus exhibited distinct peaks in concentration following the onset of the encoding (∼1.08 s) and recall periods (∼1.42 s), with both WM phases showing significant increases in concentration of Glu compared to a control condition that accounted for visual and motor demands, indicating a direct link to cognitive processes involved in WM. Importantly, GABA modulation during encoding predicted task accuracy, with greater increases in GABA concentration associated with better performance. Together, these findings provide highly temporally precise evidence of phase-specific excitatory and inhibitory dynamics during WM.

Our reconstructed time courses of Glu concentration during a cognitive task are the most comprehensive to date, with a temporal resolution of ∼0.3 s spanning 0-2.5 s post-stimulus. They show a rise in Glu, peaking 1.08 s after stimulus presentation during encoding and 1.42 s after probe presentation during recall and returning to baseline within the 2.5 s. These dynamics extend beyond previous work which identified peaks in concentration in the 0.3–1.0 s range ^21,24^. Based on this, for cognitive processes in MFC our results would recommend setting the acquisition delay to ∼1.0-1.5 s post-stimulus. Our results suggest that prior work investigating cognitive processes ^6,22,23^ may have sampled Glu too early. Indeed, we did not find a significant increase within 0-0.67 s.

Interestingly, the Glu peak at 1.42 s during recall aligns with reaction times during memory trials (1.38 s), however increases in Glu cannot solely be attributed to motor responses since no response was required during encoding. The close alignment between Glu peak and RT could reflect processes following memory retrieval, for instance internal monitoring processes which have been associated with activity in the posterior MFC ^30^. The time of the Glu peak during encoding (1.08s) corresponds to the later end of a previously hypothesized Glutamatergic Response Function (GRF) peaking 0.3-1.0 s after stimulus onset ^19^ and later than the peak of 0.5 s suggested by Koolschijn et al ^17^.

The apparent return towards baseline of the Glu response within 2.5 s during encoding potentially suggests a transient response which may not reflect maintenance. Maintenance may be served by the dlPFC instead ^32–34^. Moreover, the return to baseline within each WM phase indicates that we could detect distinguishable responses to each phase, rather than a single prolonged response across the WM block.

In our analysis of longer time windows, windowing improved the visibility of the Glu response. When incorporating all sampling points, we observed some evidence of increased Glu during encoding (1.4%) and recall (1.8%) compared to control, significant prior to Bonferroni correction. As expected from the observed time course, windowing to the central eight sampling points (0.67-1.83 s), thereby focussing on the peaks of the Glu responses, revealed larger detectable increases in Glu during encoding (2.5%) and recall (3.2%). The changes in Glu averaged across this window are less than previous work in the hippocampus, which similarly compared encoding and recall to a visuo-motor control task (4-10%) ^35^ and in the visual cortex when comparing remembered to forgotten items (∼5%) ^18^. However, similar magnitude changes were reported in the dlPFC during a WM task (2.7%) ^16^. Over our measured time courses, we find maximal percent Glu changes of 5.4% and 5.3% during encoding and retrieval, respectively, relative to control. We may have observed larger % increases had we performed a more classic event-related design, intermixing control and WM trials, rather than a mixed-design. While we observed a rise and fall of Glu over the 2.5 window, the magnitude of this transient response may have been partially hidden by slower changes across the block of trials, speculated to be involved in homeostatic changes or plasticity ^36^.

The increase in Glu during WM trials compared to control trials is consistent with the increased fMRI activation for working memory over control trials in our fMRI localiser (see Methods and Supplementary Methods). In contrast, Jelen and colleagues ^7^ did not find a main effect of working memory load (0-back versus 2-back) on Glx (Glu and glutamine) in the MFC, however, here we employed a different type of WM task. We also acquired fMRS at 7T, which offers a larger signal-to-noise ratio (SNR) and improved sensitivity to separate signals such as Gln and Glu ^37^.

Interestingly, individual increases in Glu did not correlate with WM performance. The increase in Glu might therefore reflect either general increases in attentional demands during the WM task, not directly tied to performance and/or it could be linked to cognitive control processes ^38^ that are not (or to a lesser extent) required in the control condition. The posterior MFC has been associated with cognitive control processes before ^e.g., 30^. The WM task used in this study involves associative binding of different colours, shapes and locations, thereby creating a representational competition and potential response conflict. Higher Glu levels during the WM task compared to the simpler control task indicate that the metabolic demand is tied to encoding and recalling complex associations, not just visual perception or motor responses. Similar observations of increased Glx (Glu + glutamine) in medial prefrontal cortex during an associative memory task involving a longer delay period have been reported ^39^. The increases in Glu with cognitive control demands are supported by the raw Glu time courses (Fig. 1a), with increases during WM encoding and recall (compared to baseline) but little change in the control task. The control task used here elicited minimal or no conflict because the response mapping was direct and stable. If increased cognitive control demands are managed successfully, as in adaptive proactive control processes ^40^ leading to a resolution of internal response conflicts before a response is executed, a direct association with behavioural task performance is not expected. The decreasing size of Glu responses over blocks (Fig. 3c) would also be in line with cognitive control processes: as participants get more practiced to execute the WM task, some processes might become more automatised (e.g. due to the development of strategies to memorise associations), thereby decreasing cognitive control demands. An increase with cognitive demands is consistent with evidence showing a positive correlation between increased Glx and BOLD (blood oxygenation level dependent) signal in the dlPFC and lower WM performance, which could reflect the increased neurovascular demands of challenging cognitive processes ^41^.

The diminishing increase in Glu from the first to the third run, could not be explained by build-up of Glu over the session as there was no difference in the baseline Glu concentration measured over subsequent runs during the initial rest period (8 spectra). The result may also be explained by repetition suppression – repeating stimuli can result in a decrease in neural response that is detectable within neuroimaging ^42^. This effect has more recently been detected in the glutamate signal using ^1^H-MRS ^21,24^. While no exact combination of shape, location, colour and probe was repeated in our study, we may have observed repetition suppression to the general type of WM trials.

In addition to its role as an excitatory neurotransmitter, Glu plays a key role in cerebral metabolism through its involvement in the tricarboxylic acid (TCA) cycle, where it can be used as a buffer. Glu is distributed intracellularly between the cytosol and synaptic vesicles, with the majority residing in the cytosol ^43^. These compartments have been suggested to exhibit distinct T₂ relaxation times, with vesicular Glu relaxing rapidly (∼10 ms) compared to cystolic Glu (180–220 ms) at 3T ^44^. Consequently, given the relatively long echo time used in our sLASER acquisition (TE = 80 ms), the observed Glu signal changes may be dominated by cystolic Glu pools rather than vesicular components. Other fMRS studies employing cognitive paradigms ^16,18,35^ have reported sustained increases in glutamate (Glu) concentration that, in some cases, are larger than those observed in the present study (2.7–10%). Notably, all these studies used short-TE MRS sequences (TE = 23–36 ms). In our work, we combined long-TE MRS with a rapidly sampled event-related paradigm in an effort to probe this faster acting synaptic component of Glu. However, we cannot separate the potential contribution of a metabolic Glu component, which may also explain the absence of a correlation with cognitive measures.

We did not observe significant group-level changes in GABA during either encoding or recall compared to control trials. Similar findings of no change in GABA levels have been reported in medial ^39^ and dorsolateral ^41^ prefrontal cortex during performance of an associative memory task. However, at an individual level, we observed that larger increases in GABA during encoding were associated with enhanced task performance, with similar trends present during recall. This suggests that dynamic, individual-level GABA fluctuations underpin aspects of WM performance.

This study is the first to show a relationship between GABA dynamics within MFC during the encoding phase of a WM task and performance. Previous work with static MRS has highlighted a positive relationship between WM performance and GABA quantified at rest from another frontal region - the dlPFC ^45,46 although see 47,48^.

Interestingly, Yoon et al ^46^ found that higher GABA levels were predictive of improved performance at higher WM loads. Higher GABA may therefore be particularly beneficial for accurate performance on demanding WM tasks such as the one used here, potentially via suppression of task irrelevant information – a role already observed in MFC through measuring the BOLD signal with fMRI ^26,27^. In MFC, GABA increases have been observed during interference tasks requiring inhibition ^49^ and higher hippocampal GABA levels have been linked to improved performance during a think/no-think paradigm in which task irrelevant information is suppressed ^50^.

Since Glu and GABA seem to be related to different types of cognitive processes (see also review by Pasanta and colleagues ^36^), it could be beneficial to investigate Glu and GABA dynamics separately, rather than combining these measures into excitatory/inhibition ratios ^e.g., see 8^ in the context of WM tasks.

To conclude, based on our findings, we can now better characterise the time course of Glu responses in relation to different events in cognitive tasks, with the change from control trials peaking around 1.08 s during encoding and 1.42 s during recall. Our results also demonstrated the benefits and importance of selecting the appropriate acquisition time to be maximally sensitive to Glu changes. Comparing WM-related metabolite changes to a control condition allowed us to isolate the changes associated with cognitive functions involved in WM over and above visual and motor responses. Future theoretical work on the roles of Glu and GABA in WM should integrate the findings that Glu levels in MFC are closely tied to WM demands but do not predict performance, whereas GABA increases during the encoding phase do predict task performance.

## 4. Methods

### 4.1. Participants

Twenty-one participants took part in the study, two were excluded due to poor data quality (see data analysis), leaving a final sample of 19 participants (9 female, 10 male; one left-handed), aged 20-37 years (mean: 28.95, SD: 5.68 years). One participant was red-green colourblind, however they could distinguish the colours used in the task sufficiently and were not a significant outlier from either their working memory performance or neurochemical levels so were retained for analysis.

No participants reported having a current neurological or psychiatric disorder, recent medication use that could influence neurochemicals of interest, or a history of significant head injury. All participants provided informed consent. The study was approved by the local ethics committee (ethical approval number F1465).

### 4.2. Task

During the fMRS scan, participants completed four runs of an associative visuospatial WM task, adapted from Belekou et al ^51^, with two blocks of memory and two blocks of control trials per run, with each block containing 16 trials (see Fig. 4). Participants completed 24 practice trials (12 memory task, 12 control task) before scanning.

**Figure 4.**
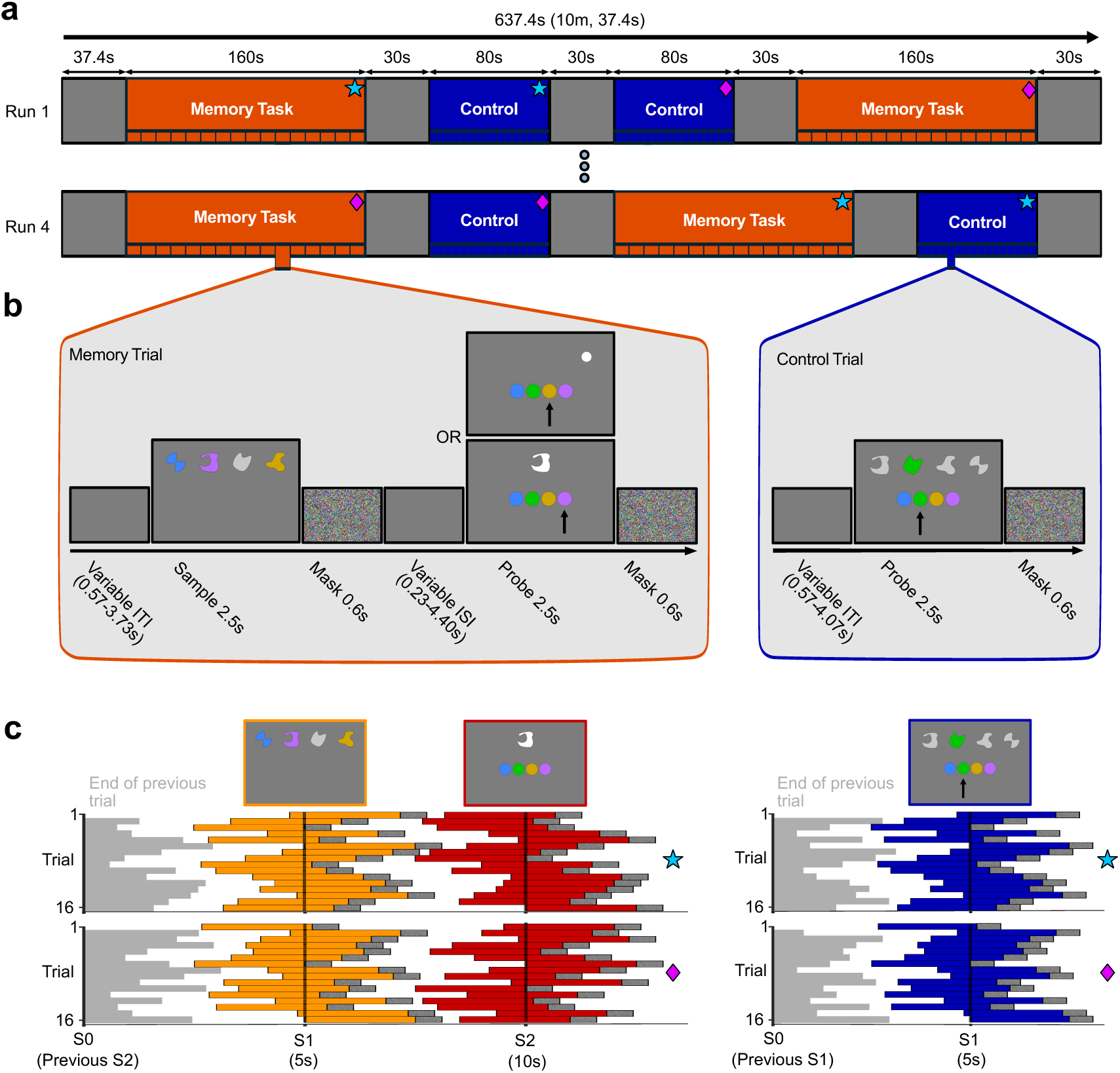
Plot of run and trial structures. *Note:* (**a**) A depiction of two runs of the task, showing two of the four counterbalancing options used in the study. Purple diamonds and red stars refer to the trial timing sets depicted in **b**. (**b**) Depictions of a memory trial (left) and control trial (right). In the memory trial, participants may be presented with a location probe (top) or shape probe (bottom). Arrows showing the correct answer are for illustration purposes only. ITI = inter-trial interval, ISI = inter-stimulus interval. (**c**) Coloured boxes show timings of stimuli in relation to sample acquisition (dashed vertical lines) for the memory trials (left) and control trials (right). The upper and lower plots show the two sets of predefined timings. Each row represents a trial within a block, starting with the top row. Textured boxes represent the mask duration. S1 = acquisition of the first MRS sample within a trial, which occurs during encoding in the memory trials, and the probe in the control trials. S2 = acquisition of the second MRS sample in the memory trials. The grey lines show when then previous trial ended.

Custom shapes without obvious semantic associations were used as stimuli in the associative working memory task. Each stimulus spanned an equal width and height and could be presented in one of four colours: blue (RGB: 62, 132, 254), green (RGB: 2, 200, 2), ochre (RGB: 208, 166, 3) or purple (RGB: 187, 106, 254).

Each WM trial started with a variable delay (grey screen) between 0.57 and 3.73 s. Participants were then shown four shapes presented at four horizontal locations spread across the width of the screen in the upper half of the display (see Fig. 4b, the encoding phase, 2.5 s). Three shapes were coloured, with participants needing to encode the shape, location and colour of the items. The fourth shape was grey, with the shape and location of the grey item varying across trials. The grey item was a placeholder for the location and did not need to be actively encoded. Participants were then presented with a scintillating, coloured mask (0.6 s) followed by a variable inter-stimulus interval (ISI). Each of the 16 trials within a block had a unique ISI ranging between 0.23 and 4.40 s. This was followed by the recall phase, in which either a white shape or location cue was presented above four response colour options (recall phase, 2.5 s). Participants had to recall the colour of the shape or the colour presented at the cued location from the encoding phase (response window: 2.5 s). The recall phase was again followed by a coloured mask (0.6 s) before the next trial began. More details about the ITIs and ISIs can be found in the data acquisition section. Memory trials spanned an average of 10 s, with a total block duration of 160 s.

The control task used the same stimuli as the associative WM task. Within a control trial participants were presented with three grey items and one coloured item above the four response options. Participants responded to the colour of the item.

The control task was designed to elicit similar visual processes and motor responses as the WM task to control for these processes in the planned analysis. This screen was presented for 2.5 s followed by a 0.6 s mask. Control trials spanned an average of 5 s including a variable ITI (0.57-4.07 s), with a total block duration of 80 s. Timings of the control trials followed the timings of the encoding phase in the memory trials.

Within each of the four runs of the task, participants were presented with two blocks of 16 memory trials and two blocks of 16 control trials, presented in a counterbalanced order. Blocks were separated by a 30 s rest period, and each run began with 37.4 s rest and ended with a 30 s rest. Each run therefore took a total of 637.4 s (10 minutes, 37.4 seconds).

The order of the trials was pseudo-randomised across participants such that no participant encountered the same trial twice, and different participants were presented with different subsets of trials. Each memory block contained an equal number of shape and location cues, and an equal number of the four options as the target colour. Timings of the trials within the block were pre-defined by two sets of possible timings (see fMRS data acquisition for more details, see also Fig. 4c).

### 4.3. Stimulus Apparatus

The task was presented using an MR compatible projector (PROPixx MRI from VPixx), which displayed the stimuli at 120 Hz on a screen suspended vertically above the foot end of the scanner bed, with participants using mirrored glasses to view the display. Screen resolution was 1980 x 1020 pixels, with the projection subtending 62 by 32 cm at approximately 2.4 m, although the lower half of the screen was generally not visible due to the participant’s body partly obstructing the field of view. The researchers ensured participants could see the stimuli before commencing the experiment. Participants responded to the task using the four fingers of their right-hand and the RESPONSEPixx 5-button response box, with each button corresponding to one of the four response options presented on the screen.

### 4.4. MR Data acquisition

Data were acquired on a 7T MR scanner (Phillips Achieva) using a 32 channel receive and 2 channel transmit head coil (Nova Medical).

An anatomical 3D T1-weighted gradient echo acquisition was first acquired and used for subsequent positioning of MRS voxel, TR = 4.9 s, voxel size = 0.85 x 0.85 x 1 mm^3^, TE = 2.17 ms, flip angle = 7°, field of view = 246 x 246 x 180 mm^3^, T_acq_ = 119 s. All single-voxel spectroscopy measurements were acquired with semi-LASER localisation and TR = 5s, voxel size = 20 x 20 x 20 mm^3^, spectral points = 2048, bandwidth = 6000 Hz, TE = 80 ms, NSA = 128. TE=80 ms was chosen to optimise sensitivity to both Glu and GABA at 7 T in line with a recent publication ^52^. Refocussing was performed using GOIA-WURST pulses ^53^. A DOTCOPS crusher gradient scheme ^54^ with interleaved outer volume suppression (OVS) and VAPOR water suppression (BW = 140 Hz) ^55^ were performed.

Voxel placement was informed by an fMRI localiser (see Supplementary Methods) captured in five separate participants using the task, with slightly modified timings and trial numbers to better suit fMRI acquisition. The MRS voxel was placed in the pre-SMA region of the MFC, parallel to but just superior to the cingulate cortex. The most posterior edge of the voxel was positioned just anterior (∼2mm) of a vertical line drawn from the AC, perpendicular to the AC-PC line, although voxel placement varied slightly per-participant to avoid larger blood vessels (see Fig. 5 for positioning).

**Figure 5.**
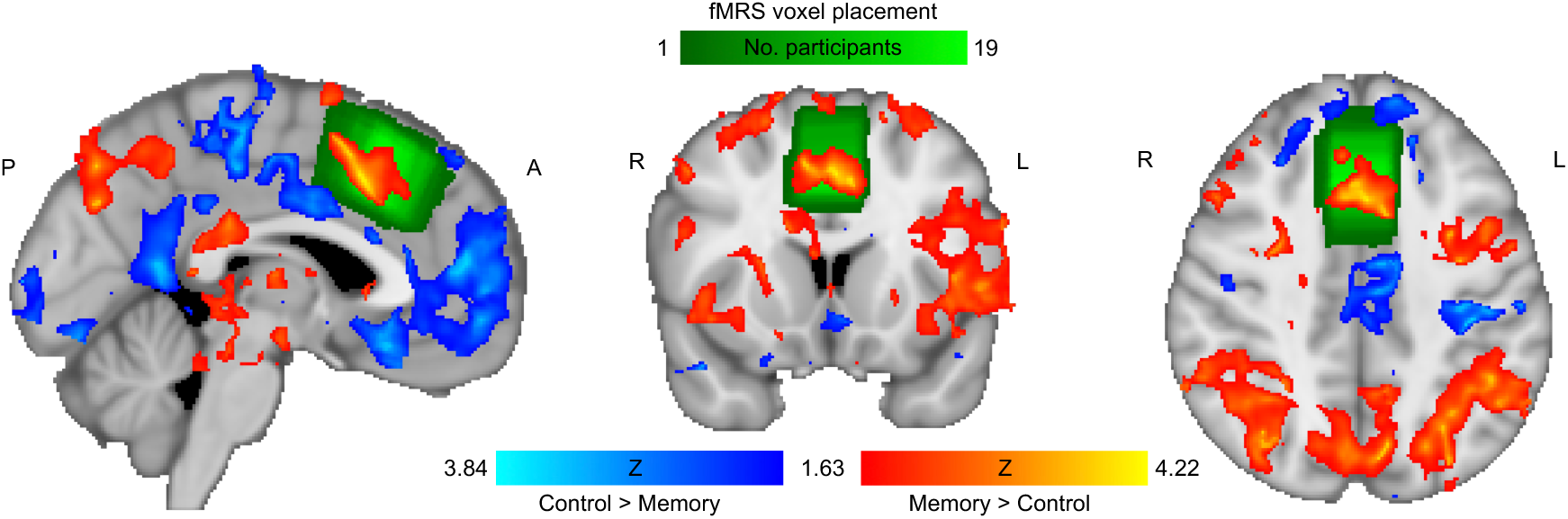
FMRI-informed positioning the fMRS voxel. *Note:* Heatmap of voxel placements over participants (green), superimposed with fMRI activation from the localiser study (see Supplementary Materials). The orange regions of activation show a greater BOLD response to memory trials over control trials. Blue regions show a greater BOLD signal to control over memory trials.

In each of the four runs, a total of 128 spectra were acquired. Within each memory trial, two spectra were acquired, one during the encoding phase and one during recall. During the control trials, a single acquisition was made per trial. Examples of spectral quality and metabolite fits can be found in Fig. 6.

**Figure 6.**
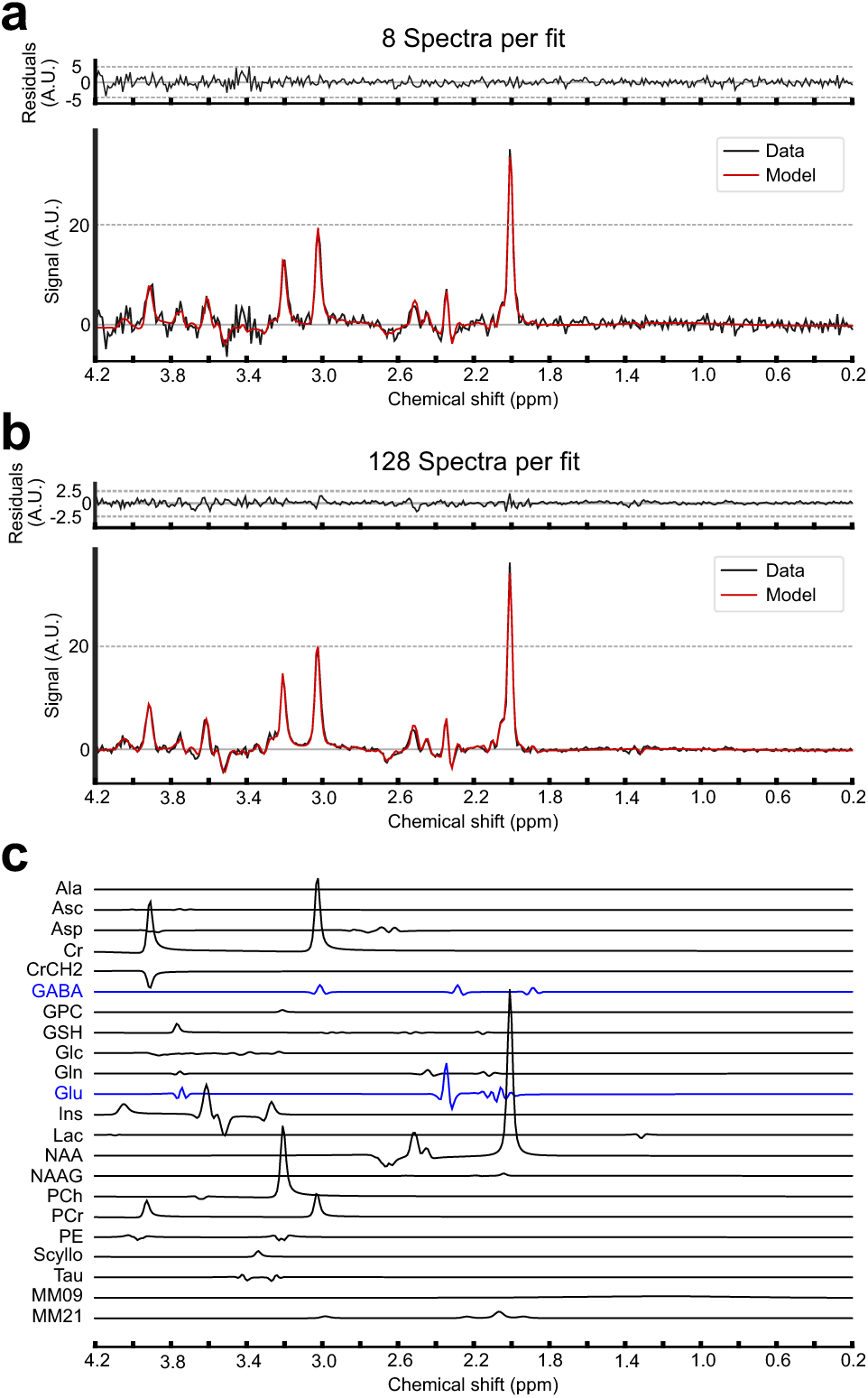
Sample spectra from one participant using 8 (**a**) and 128 (**b**) averages per fit, and composition of neurochemicals (**c**). *Note:* Example spectra acquired from a single subject, averaging the initial 8 spectra in the first rest period (**a**), and averaging over all 128 spectra acquired in a single run (**b**). All spectra have been pre-processed (black) and then fit (red). (**c**) shows the individual fits for each metabolite for included in the analysis of (b) with GABA and Glu highlighted in blue.

The onsets of the stimuli were varied such that across the 16 trials in a block, the spectral acquisitions aligned with 16 equally spaced intervals between stimulus onset (0 s) and stimulus offset (2.5 s); every 0.17 s, acquired in a non-sequential order. The timings were pre-defined, with two unique solutions designed to be maximally different (see Fig. 4c and Supplementary Materials). The sampling method of the spectral acquisitions was designed and applied to acquire neurochemical responses at different time lags after stimulus onset and to be able to estimate the glutamate response function associated with this task. The same order of sampling times from the encoding phase was used for the control trials as well (see Fig. 4c). Within a run, one memory and one control block followed each of the two solutions, with the order of the solutions being counterbalanced across runs.

### 4.5. fMRS data analysis

All spectral processing was performed in FSL-MRS V2.4.0 ^56^ in Python V3.12.7. Each run was acquired as separate scan with corresponding raw data saved for processing. Pre-processing of each run involved coil combination, eddy current-correction, frequency and phase alignments, Hankel Lanczos Singular Value Decomposition (HLSVD) and zero-order automatic phase corrections. The frequency and phase alignments were performed around the NAA singlet (1.7-2.3 ppm). Water reference spectra were used for pre-processing steps and also fit to obtain linewidth (mean: 8, SD: 3 Hz). Spectral data with water reference linewidths >20 Hz were rejected (one participant).

Fitting was performed using a truncated-Newton algorithm. Basis sets were generated using FSL-MRS via density matrix simulations using a sLASER sequence with real pulse information and interpulse timings. with TE=80 ms, and a 90° spredrex and 180° adiabatic GOIA pulses. The metabolites generated were Alanine (Ala), Ascorbate (Asc), Aspartate (Asp), Creatine (Cr), Creatine CH2 group (CrCH2), Gamma-aminobutyric acid (GABA), Glucose (Glc), Glutamine (Gln), Glutamate (Glu), Glycine (Gly), Glycerophosphocholine (GPC), Glutathione (GSH), Myo-Inositol (Ins), Lactate (Lac), N-acetylaspartate (NAA), N-acetylaspartylglutamate (NAAG), Phosphocholine (PCh), Phosphocreatine (PCr), Phosphoethanolamine (PE), Scyllo-Inositol (Scyllo) and Tau. No baseline was used and simulated macromolecular components at 0.9 and 2.1 ppm were implemented. Each macromolecule was assigned to a separate group for linewidth and phase compared to the rest of the other metabolites. To form a high SNR fit during initialisation for further analysis, all spectra from the four runs per participant are averaged together and fit together, an appropriate scaling is obtained from this data as well per participant

#### 4.5.1. Time course analysis

The time course analysis was achieved by assigning spectra for the 16 sampling points into to eight bins (or timepoints), with the mean sampling time for each bin as follows: 0.083, 0.417, 0.75, 1.083, 1.417, 1.750, 2.083, 2.417 s. For each bin, that resulted in a total of four spectral acquisitions per run. The time courses were initially converted to a percentage change compared to the initial rest spectra for each run and then averaged across runs, to observe the dynamic change in Glu during each WM phase (encoding, recall) and during the control trials. One-sample t-tests were used to assess whether the concentration during each of the WM phases and control trials was significantly different from the initial rest period (baseline) at any timepoint. Bonferroni-corrected α = .002.

For each timepoint and run separately, the Glu concentrations during encoding and recall were additionally calculated as a percentage change from the control condition, without any normalisation to the initial rest spectra, to isolate processes involved in WM. Percent changes were then averaged over run separately for each participant. One-sample t-tests were again employed to assess if Glu levels during encoding and recall were significantly different at any timepoint compared to control. T-tests were two-sided. Bonferroni-corrected α = .003.

#### 4.5.2. WM phase analysis

In addition to the time course analysis, we performed event-based analysis averaging over larger time windows. Spectra for each run and task phase (control, encoding, recall) separately were averaged over either all 16 sampling points (number of spectral averages, NSA=32/run/phase), the middle eight (NSA=16/run/phase), or the first and last four sampling points (NSA=16/run/phase, see Results) prior to fitting. The change in Glu and GABA concentrations were calculated as the percentage change during encoding and recall compared to the control task to capture memory-related changes in concentrations over the visual and motor responses in the control trials. No internal, within-spectra normalisation, such as to total creatine, was performed. The normalisation to control was performed for each run separately and then averaged over runs to assess for effects of WM phase, or averaged over encoding and recall assessing the main effect of run.

We assessed whether 1) the concentration changes during encoding and recall differed from control trials (one-sample t-tests), 2) the concentration changes differed between encoding and recall (paired-sample t-tests), 3) the concentration changes were significantly correlated across encoding and recall (Pearson’s correlation). T-tests were two-sided.

GABA levels for one participant were removed due to being a significant outlier during recall (>3 SD from group mean).

### 4.6. Behavioural data analysis

All trials were retained in the behavioural data analysis as no trials showed a pre-emptive response (<0.3 s). Of 256 trials, including both memory and control trials, a maximum of eight trials were missed by a single participant. Missed trials in a WM task may reflect a lapse in attention or may reflect subjective task difficulty. RT-based outliers in individual trials were negligible (max. 5 trials for an individual participant). Including trial wise outliers should therefore not compromise data quality.

To compare accuracy and reaction time over the four runs, within-subjects one-way ANOVAs and post-hoc paired samples t-tests were employed. T-tests were two-sided.

## Supporting information

MRSinMRS

## 5.#Acknowledgements

This work was supported by the Wellcome Trust [226715/Z/22/Z]. AB would like to acknowledge the Royal Academy of Engineering. We thank Prof. Helen Barron for their insight into study design during the early stages of the project, Dr. Josh Khoo for creating an initial version of the experimental script, and Siavash Rakhtshah for assisting with MR scanning and with eligibility queries.

## 7. Author contributions

All authors were involved in task-design. AB and DC optimised fMRS data acquisition and analysis. RE created the experimental script. DC and RE collected and analysed the data with supervision from AB, CD and KD. RE, DC, CD, KD and AB wrote the manuscript. All authors reviewed and approved the manuscript. DC and RE have contributed equally to the manuscript and therefore share first authorship.

## References

1. Patel, T. & Talcott, J. B. Moderate relationships between NAA and cognitive ability in healthy adults: implications for cognitive spectroscopy. Front. Hum. Neurosci. 8, (2014).

2. Reid, M. A. et al. 7T Proton Magnetic Resonance Spectroscopy of the Anterior Cingulate Cortex in First-Episode Schizophrenia. Schizophr. Bull. 45, 180–189 (2019).

3. Rowland, L. M. et al. Frontal Glutamate and γ-Aminobutyric Acid Levels and Their Associations With Mismatch Negativity and Digit Sequencing Task Performance in Schizophrenia. JAMA Psychiatry 73, 166–174 (2016).

4. White, T. L. et al. The neurobiology of wellness: 1H-MRS correlates of agency, flexibility and neuroaffective reserves in healthy young adults. NeuroImage 225, 117509 (2021).

5. Jelen, L. A., King, S., Mullins, P. G. & Stone, J. M. Beyond static measures: A review of functional magnetic resonance spectroscopy and its potential to investigate dynamic glutamatergic abnormalities in schizophrenia. J. Psychopharmacol. (Oxf.) 32, 497–508 (2018).

6. Craven, A. R. et al. GABA, glutamatergic dynamics and BOLD contrast assessed concurrently using functional MRS during a cognitive task. NMR Biomed. 37, e5065 (2024).

7. Jelen, L. A. et al. Functional magnetic resonance spectroscopy in patients with schizophrenia and bipolar affective disorder: Glutamate dynamics in the anterior cingulate cortex during a working memory task. Eur. Neuropsychopharmacol. 29, 222–234 (2019).

8. Saviola, F., et al. Disentangling metabolic and neurovascular timescales supporting cognitive processes. Proc. Natl. Acad. Sci. 122, e2506513122 (2025).

9. Bednařík, P. et al. Neurochemical responses to chromatic and achromatic stimuli in the human visual cortex. J. Cereb. Blood Flow Metab. 38, 347–359 (2018).

10. Chen, C. et al. Activation induced changes in GABA: Functional MRS at 7 T with MEGA-sLASER. NeuroImage 156, 207–213 (2017).

11. DiNuzzo, M. et al. Perception is associated with the brain’s metabolic response to sensory stimulation. eLife 11, e71016 (2022).

12. Ip, I. B. et al. Combined fMRI-MRS acquires simultaneous glutamate and BOLD-fMRI signals in the human brain. NeuroImage 155, 113–119 (2017).

13. Mangia, S. et al. Sustained Neuronal Activation Raises Oxidative Metabolism to a New Steady-State Level: Evidence from 1H NMR Spectroscopy in the Human Visual Cortex. J. Cereb. Blood Flow Metab. 27, 1055–1063 (2007).

14. Mekle, R. et al. Detection of metabolite changes in response to a varying visual stimulation paradigm using short-TE 1H MRS at 7 T. NMR Biomed. 30, e3672 (2017).

15. Volovyk, O. & Tal, A. Increased Glutamate concentrations during prolonged motor activation as measured using functional Magnetic Resonance Spectroscopy at 3T. NeuroImage 223, 117338 (2020).

16. Woodcock, E. A., Anand, C., Khatib, D., Diwadkar, V. A. & Stanley, J. A. Working Memory Modulates Glutamate Levels in the Dorsolateral Prefrontal Cortex during 1H fMRS. Front. Psychiatry 9, (2018).

17. Koolschijn, R. S., Clarke, W. T., Ip, I. B., Emir, U. E. & Barron, H. C. Event-related functional magnetic resonance spectroscopy. NeuroImage 276, 120194 (2023).

18. Koolschijn, R. S. et al. Memory recall involves a transient break in excitatory-inhibitory balance. eLife 10, e70071 (2021).

19. Mullins, P. G. Towards a theory of functional magnetic resonance spectroscopy (fMRS): A meta-analysis and discussion of using MRS to measure changes in neurotransmitters in real time. Scand. J. Psychol. 59, 91–103 (2018).

20. Lea-Carnall, C., El-Deredy, W., Stagg, C. J., Williams, S. R. & Trujillo-Barreto, N. A mean-field model of glutamate and GABA synaptic dynamics for functional MRS. NeuroImage 266, 1–18 (2023).

21. Apšvalka, D., Gadie, A., Clemence, M. & Mullins, P. G. Event-related dynamics of glutamate and BOLD effects measured using functional magnetic resonance spectroscopy (fMRS) at 3 T in a repetition suppression paradigm. NeuroImage 118, 292–300 (2015).

22. Craven, A. R. et al. GABA, glutamate dynamics and BOLD observed during cognitive processing in psychosis patients with hallucinatory traits. Sci. Rep. 15, 19466 (2025).

23. Mohammadi, H., et al. Unveiling Glutamate Dynamics: Cognitive Demands in Human Short-Term Memory Learning Across Frontal and Parieto-Occipital Cortex: A Functional MRS Study. J. Biomed. Phys. Eng. 14, 519–532 (2024).

24. Lally, N. et al. Glutamatergic correlates of gamma-band oscillatory activity during cognition: A concurrent ER-MRS and EEG study. NeuroImage 85, 823–833 (2014).

25. Baddeley, A. D. & Hitch, G. Working Memory. in Psychology of Learning and Motivation (ed. Bower, G. H.) vol. 8 47–89 (Academic Press, 1974).

26. Danielmeier, C., Eichele, T., Forstmann, B. U., Tittgemeyer, M. & Ullsperger, M. Posterior Medial Frontal Cortex Activity Predicts Post-Error Adaptations in Task-Related Visual and Motor Areas. J. Neurosci. 31, 1780–1789 (2011).

27. Danielmeier, C. et al. Acetylcholine Mediates Behavioral and Neural Post-Error Control. Curr. Biol. 25, 1461–1468 (2015).

28. Klein, T. A. et al. Neural correlates of error awareness. NeuroImage 34, 1774–1781 (2007).

29. Rushworth, M. F., Buckley, M. J., Behrens, T. E., Walton, M. E. & Bannerman, D. M. Functional organization of the medial frontal cortex. Curr. Opin. Neurobiol. 17, 220–227 (2007).

30. Ullsperger, M., Danielmeier, C. & Jocham, G. Neurophysiology of Performance Monitoring and Adaptive Behavior. Physiol. Rev. 94, 35–79 (2014).

31. Kamiński, J. et al. Persistently active neurons in human medial frontal and medial temporal lobe support working memory. Nat. Neurosci. 20, 590–601 (2017).

32. Barbey, A. K., Koenigs, M. & Grafman, J. Dorsolateral prefrontal contributions to human working memory. Cortex 49, 1195–1205 (2013).

33. Goldman-Rakic, P. S. Circuitry of the frontal association cortex and its relevance to dementia. Arch. Gerontol. Geriatr. 6, 299–309 (1987).

34. Goldman-Rakic, P. S. Regional and cellular fractionation of working memory. Proc. Natl. Acad. Sci. 93, 13473–13480 (1996).

35. Stanley, J. A. et al. Functional dynamics of hippocampal glutamate during associative learning assessed with in vivo 1H functional magnetic resonance spectroscopy. NeuroImage 153, 189–197 (2017).

36. Pasanta, D. et al. Functional MRS studies of GABA and glutamate/Glx – A systematic review and meta-analysis. Neurosci. Biobehav. Rev. 144, 104940 (2023).

37. Tkáč, I., Öz, G., Adriany, G., Uğurbil, K. & Gruetter, R. In vivo 1H NMR spectroscopy of the human brain at high magnetic fields: Metabolite quantification at 4T vs. 7T. Magn. Reson. Med. 62, 868–879 (2009).

38. Botvinick, M. M., Braver, T. S., Barch, D. M., Carter, C. S. & Cohen, J. D. Conflict monitoring and cognitive control. Psychol. Rev. 108, 624–652 (2001).

39. Thielen, J.-W. et al. The increase in medial prefrontal glutamate/glutamine concentration during memory encoding is associated with better memory performance and stronger functional connectivity in the human medial prefrontal–thalamus–hippocampus network. Hum. Brain Mapp. 39, 2381–2390 (2018).

40. Braver, T. S. The variable nature of cognitive control: a dual mechanisms framework. Trends Cogn. Sci. 16, 106–113 (2012).

41. Oh, H. et al. A preliminary study of dynamic neurochemical changes in the dorsolateral prefrontal cortex during working memory. Eur. J. Neurosci. 59, 2075–2086 (2024).

42. Grill-Spector, K. & Malach, R. fMR-adaptation: a tool for studying the functional properties of human cortical neurons. Acta Psychol. (Amst.) 107, 293–321 (2001).

43. Walker, M. C. & van der Donk, W. A. The many roles of glutamate in metabolism. J. Ind. Microbiol. Biotechnol. 43, 419–430 (2016).

44. Emeliyanova, P., Parkes, L. M., Williams, S. R. & Lea-Carnall, C. Evidence for biexponential glutamate T2 relaxation in human visual cortex at 3T: A functional MRS study. NMR Biomed. 37, e5240 (2024).

45. Ragland, J. D. et al. Disrupted GABAergic facilitation of working memory performance in people with schizophrenia. NeuroImage Clin. 25, 102127 (2020).

46. Yoon, J. H., Grandelis, A. & Maddock, R. J. Dorsolateral Prefrontal Cortex GABA Concentration in Humans Predicts Working Memory Load Processing Capacity. J. Neurosci. 36, 11788–11794 (2016).

47. Andersson, P., Li, X. & Persson, J. Hippocampal and prefrontal GABA and glutamate concentration contribute to component processes of working memory in aging. Cereb. Cortex 35, bhaf105 (2025).

48. Michels, L. et al. Frontal GABA Levels Change during Working Memory. PLOS ONE 7, e31933 (2012).

49. Kühn, S. et al. Neurotransmitter changes during interference task in anterior cingulate cortex: evidence from fMRI-guided functional MRS at 3 T. Brain Struct. Funct. 221, 2541–2551 (2016).

50. Schmitz, T. W., Correia, M. M., Ferreira, C. S., Prescot, A. P. & Anderson, M. C. Hippocampal GABA enables inhibitory control over unwanted thoughts. Nat. Commun. 8, 1311 (2017).

51. Belekou, A., Katshu, M. Z. U. H., Dundon, N. M., d’Avossa, G. & Smyrnis, N. Spatial and non-spatial feature binding impairments in visual working memory in schizophrenia. Schizophr. Res. Cogn. 32, 100281 (2023).

52. Finkelman, T., Furman-Haran, E., Paz, R. & Tal, A. Quantifying the excitatory-inhibitory balance: A comparison of SemiLASER and MEGA-SemiLASER for simultaneously measuring GABA and glutamate at 7T. NeuroImage 247, 118810 (2022).

53. Tannús, A. & Garwood, M. Improved Performance of Frequency-Swept Pulses Using Offset-Independent Adiabaticity. J. Magn. Reson. 120, 133–137 (1996).

54. Landheer, K. & Juchem, C. Dephasing optimization through coherence order pathway selection (DOTCOPS) for improved crusher schemes in MR spectroscopy. Magn. Reson. Med. 81, 2209–2222 (2019).

55. Tkáč, I., Starčuk, Z., Choi, I.-Y. & Gruetter, R. In vivo 1H NMR spectroscopy of rat brain at 1 ms echo time. Magn. Reson. Med. 41, 649–656 (1999).

56. Clarke, W. T., Ligneul, C., Cottaar, M., Ip, I. B. & Jbabdi, S. Universal dynamic fitting of magnetic resonance spectroscopy. Magn. Reson. Med. 91, 2229–2246 (2024).

