## Supplementary material for "Temporally resolved glutamate and GABA responses measured in human medial frontal cortex during working memory encoding and recall": MRSinMRS

**Table 1**. MRSinMRS checklist. Additional columns are provided for multi-site or multi-sequence studies if necessary.

| Site (Name or Number)<br><br>1. Hardware | |
|---|---|
| a. Field strength [T] | 7T |
| b. Manufacturer | Philips |
| c. Model (software version if available) | Achieva |
| d. RF coils: nuclei (transmit/ receive), number of channels, type, body part | Nova Medical T/R (2/32) head coil |
| e. Additional hardware | MR compatible projector PROPixx MRI from VPixx<br><br>RESPONSEPixx 5-button response box |
| 2. Acquisition | |
| a. Pulse sequence | Single-voxel sLASER |
| b. Volume of Interest (VOI) locations | Media Frontal Cortex (MFC) |
| c. Nominal VOI size [$cm^3$, $mm^3$] | 20 x 20 x 20 $mm^3$ |
| d. Repetition Time (TR), Echo Time (TE) [ms, s] | TR: 5s, TE: 80 ms |
| e. Total number of Excitations or acquisitions per spectrum<br><br>In time series for kinetic studies<br><br>i. Number of Averaged spectra (NA) per time-point<br>ii. Averaging method (e.g. block-wise or moving average)<br>iii. Total number of spectra (acquired / in time-series) | Block-wise averaged spectra<br><br>Phase Analysis: 32 spectra averaged together depending on whether they were acquired during control, encoding or recall.<br><br>Timecourse Analysis: 4 spectra averaged together depending on whether they were acquired during control, encoding and recall and what the stimulus onset time is. Such that there are 8 timepoint bins per phase (control, encoding and recall). |
| f. Additional sequence parameters<br><br>(spectral width in Hz, number of spectral points, frequency offsets)<br><br>If STEAM:, Mixing Time (TM)<br><br>If MRSI: 2D or 3D, FOV in all directions, matrix size, acceleration factors, sampling method | BW: 6000 Hz<br><br>Spectral Points: 2048<br><br>GOIA RF pulses<br><br>Outer Volume Suppression<br><br>DOTCOPS Gradient crushing scheme |

| | |
|---|---|
| g. Water Suppression Method | VAPOR (Bandwidth: 140 Hz) |
| h. Shimming Method, reference peak, and thresholds for "acceptance of shim" chosen | Pencil-Beam 2$^{nd}$ order shimming |
| i. Triggering or motion correction method<br><br>(respiratory, peripheral, cardiac triggering, incl. device used and delays) | Functional task triggered by beginning of spectroscopy. |
| 3. Data analysis methods and outputs | |
| a. Analysis software | FSL-MRS V2.4.0<br><br>Python V3.12.7 |
| b. Processing steps deviating from quoted reference or product | Pre-Fitting: coil combination, eddy current-correction, frequency and phase alignments, Hankel Lanczos Singular Value Decomposition (HLSVD) and zero-order automatic phase correction.<br><br>Fitting: Truncated Newton Algorithm.<br><br>Basis Metabolites included: Alanine (Ala), Ascorbate (Asc), Aspartate (Asp), Creatine (Cr), Creatine CH2 group (CrCH2), Gamma-aminobutyric acid (GABA), Glucose (Glc), Glutamine (Gln), Glutamate (Glu), Glycine (Gly), Glycerophosphocholine (GPC), Glutathione (GSH), Myo-Inositol (Ins), Lactate (Lac), N-acetylaspartate (NAA), N-acetylaspartylglutamate (NAAG), Phosphocholine (PCh), Phosphocreatine (PCr), Phosphoethanolamine (PE), Scyllo-Inositol (Scyllo) and Tau. Simulated macromolecules at 0.9and 2.1 ppm. Ge<br><br>Basis sets were generated using FSL-MRS via density matrix simulations using a sLASER sequence with TE=80 ms, and a 90° spredrex and 180° adiabatic GOIA pulses. Using 1.8567 x $10^{-3}$, 1.3607 x $10^{-3}$, 1.7276 x $10^{-3}$, 29.6409 x $10^{-3}$ and 26.6147 x $10^{-3}$ ms delays and a rephase area following the initial 90 degree pulse in the z-direction of -2.86 x $10^{-3}$ $sm^{-1}$ |
| c. Output measure<br><br>(e.g. absolute concentration, institutional units, ratio) | Raw fitted values relative to control period or to initial rest period, reported as a percentage. |
| d. Quantification references and assumptions, | Voigt lineshape. |

| | |
|---|---|
| fitting model assumptions | |
| 4. Data Quality | |
| a. Reported variables<br><br>(SNR, Linewidth (with reference peaks)) | Mean Fit Water Linewidth: 8 ± 3 Hz. |
| b. Data exclusion criteria | Water linewidth > 20 Hz |
| c. Quality measures of postprocessing Model fitting (e.g. CRLB, goodness of fit, SD of residual) | Glu CRLB of phase analysis: 5.7 ± 0.4 %<br><br>GABA CRLB of phase analysis: 21.48 ± 1.40% (Control), 23.01 ± 1.60% (Encoding), and 22.52 ± 1.29% (Recall).<br><br>Glu CRLB of timecourse analysis: 12 ± 1 %. |
| d. Sample Spectrum | See Fig. 6 in the main manuscript. |
