## Supplementary Materials for "Temporally resolved glutamate and GABA responses measured in human medial frontal cortex during working memory encoding and recall"

#### 1. Supplementary Results

##### 1.1. Glu did not differ in the 'outside' time window

In the 'outside' time points (the first four and last four sampling points), Glu levels did not differ during encoding (mean [SE] = -0.20% [0.95%], $t(18) = -0.21$, $p = .839$, 95%CI = [-2.19, 1.80], $d = 0.05$) or recall (mean [SE] = 0.64% [1.00%], $t(18) = 0.64$, $p = .528$, 95%CI = [-1.45, 2.74], $d = 0.15$) compared to during control trials. The concentrations changes during encoding and recall were significantly correlated ($r = 0.51$, $p = .024$). See Fig. 1 in the paper. CLRBs did not significantly differ between control, encoding and recall phases using the 'outside' sampling points: 7.12% (0.66%), 6.95% (0.56%) and 7.04% (0.63%) respectively. They did also not differ from the CRLBs using the 'peak' window.

##### 1.2. GABA did not differ with WM phase across the full 2.5 s window

Averaged across the 16 sampling points, GABA did not significantly differ from control trials during encoding (mean [SE] = 0.48% [2.23%], $t(18) = 0.21$, $p = .834$, 95%CI = [-4.22, 5.17], $d = 0.05$) or recall (mean [SE] = 1.91% [2.15%], $t(18) = 0.88$, $p = .388$, 95%CI = [-2,62, 6.43], $d = 0.20$). The concentrations during encoding and recall were not significant different ($t(18) = 0.94$, $p = .362$, 95%CI = [-1.78, 4.64], $d = 0.21$) and were highly significantly correlated ($r = 0.76$, $p < .001$). See Supplementary Fig. 1. GABA CLRBs did not significantly differ between control, encoding and recall phases: 18.01% (1.07%), 18.52% (1.18%) and 18.19% (1.18%) respectively.

##### 1.3. GABA was slightly but not significantly elevated in the 'outside' time window

For analysing the outside eight sampling points, we removed the participant defined as an outlier from the middle eight time points for consistency. GABA was increased but did not significantly differ from control trials during encoding (mean

[SE] = 4.73% [2.77%], $t(17) = 1.71$, $p = .106$, 95%CI = [-1.11, 10.58], $d = 0.40$) or recall (mean [SE] = 5.01% [3.38%], $t(17) = 1.48$, $p = .156$, 95%CI = [-2.12, 12.14], $d = 0.10$). The concentrations did not significantly differ between encoding and recall ($t(17) = 0.13$, $p = .897$, 95%CI = [-4.21, 4.77], $d = 0.03$) and were highly significantly correlated ($r = 0.78$, $p < .001$). See Supplementary Fig. 1. GABA CLRBs did not significantly differ between control, encoding and recall phases: 22.36% (1.53%), 21.52% (1.39%) and 22.35% (2.00%) respectively. They also did not significantly differ from the CRLBs during the middle time window.

Supplementary Figure 1

*GABA concentrations separated by 'middle' and 'outside' time windows.*

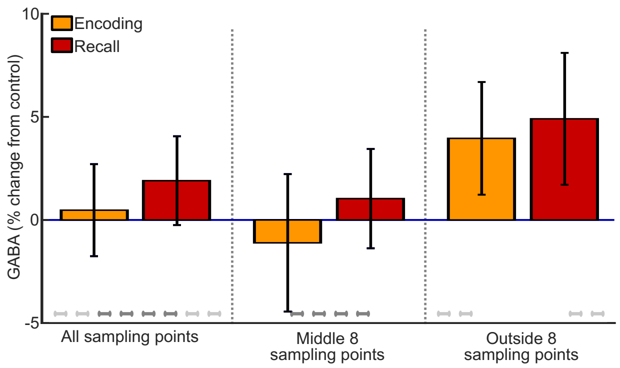


*Note:* Average % changes in GABA compared to control trials during encoding (orange) and recall (red) across all 16 sampling points (left), the middle eight sampling points (middle) and the outside eight sampling pints (right). All error bars show the standard error across participants.

#### 1.4. GABA during encoding predicts WM accuracy with the full time window

When including all 16 sampling timepoints, the change in GABA from control trials during encoding was marginally but significantly positively correlated with working memory accuracy ($r = 0.51$, $p = .025$). There was also a positive trend between GABA change during recall with accuracy ($r = 0.40$, $p = .086$) and a negative trend with reaction time ($r = -0.44$, $p = .059$) prior to correction. Bonferroni-corrected $\alpha = .025$. See Supplementary Fig. 2b.

Supplementary Figure 2

Correlations of Glu and GABA changes with WM performance using the full time window

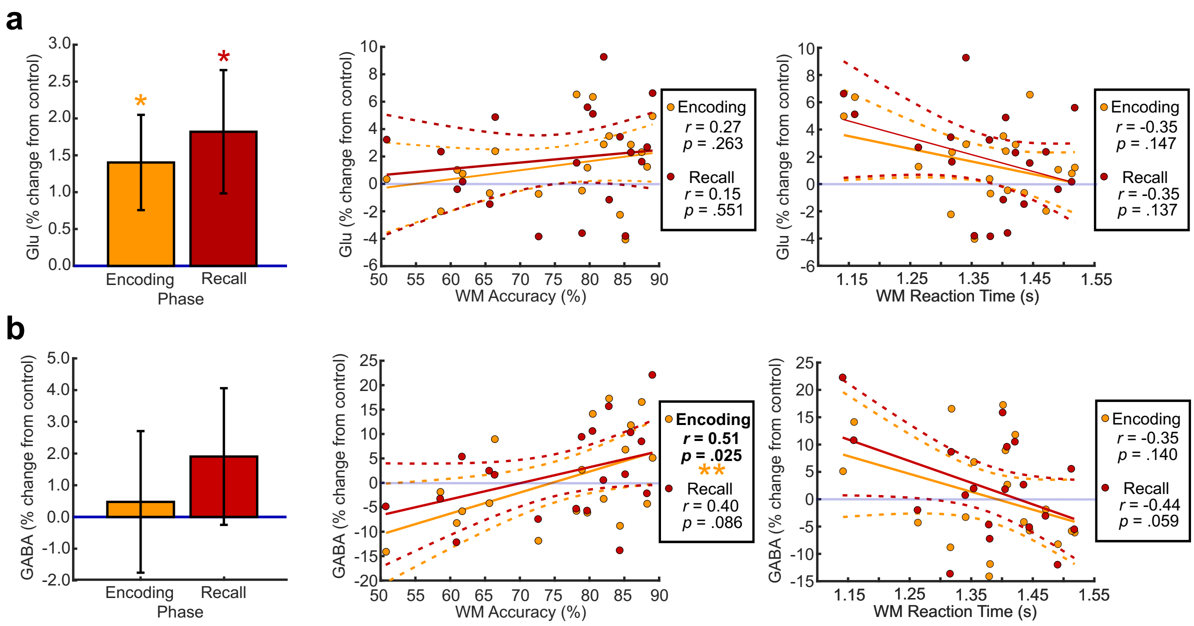


*Note:* Percent signal change from control trials during encoding (orange) and recall (red), showing changes in Glu (**a**) and GABA (**b**), showing the group-level changes (left) and the correlations with working memory accuracy (middle) and working memory reaction time (right). Plots show data using all 16 sampling points.

#### 1.5. Glu and GABA across runs with the full time window

For Glu using all 16 sampling points, mean percent signal changes (SE) per run were: Run 1: 3.30% (1.01%), Run 2: 2.47% (1.26%), Run 3: 0.23% (1.02%), Run 4: 0.45% (0.86%). A within-subjects ANOVA showed that the effect of run trended towards significance ($F(3, 54) = 2.51$, $p = .069$, $\eta_p^2 = 0.12$). Prior to correction there was a significant decrease in the Glu change in Run 3 compared to Run 1 ($t(18) = -2.12$, $p = .048$, 95% CI = [-6.11, -0.02], $d = 0.49$). The differences between Run 1 and Run 4 ($p = .060$) and between Run 2 and Run 3 ($p = .074$) only trended towards significance prior to correction. Bonferroni-corrected $\alpha = .008$. See Supplementary Fig. 3a.

One-sample t-tests compared to 0 showed a significant increase in Glu compared to control trials for Run 1 only ($t(18) = 3.25$, $p = .004$, 95% CI = [1.17, 5.43], $d = 0.75$). The increase in Run 2 only trended towards significance prior to correction ($t(18) = 1.97$, $p = .065$, 95% CI = [-0.17, 5.11], $d = 0.45$). Bonferroni-corrected $\alpha = .013$.

For GABA, the effect of run was not significant using all 16 sampling points ($F(3, 54) = 0.50$, $p = .683$, $\eta_p^2 = 0.03$). The mean percent changes (SE) were: Run 1: 4.31% (4.53%), Run 2: -2.14% (3.34%), Run 3: 1.98% (4.27%), Run 4: 0.61% (3.23%).

Supplementary Figure 3

Glu and GABA changes across runs

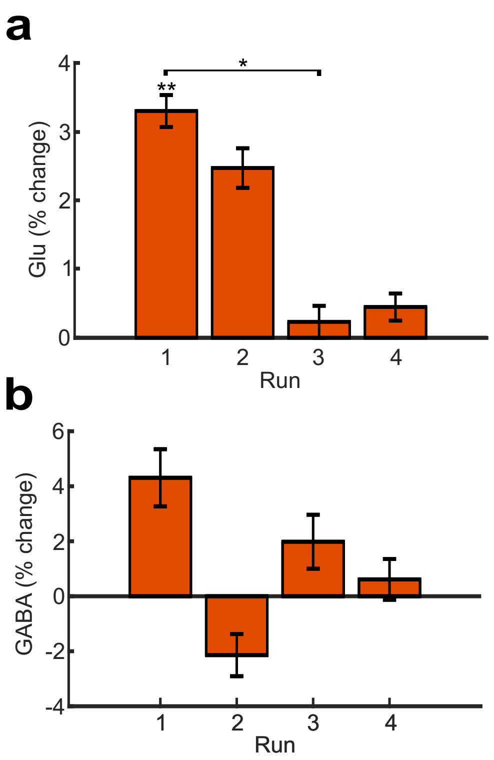


*Note:* Percent changes in Glu (**a**) and GABA (**b**) across runs compared to control trials, averaged over encoding and recall using the full time window with all 16 sampling points.

### 2. Supplementary Methods

#### 2.1. fMRI localiser

An fMRI localiser experiment was conducted to inform voxel placement.

##### 2.1.1. Participants

Five participants took part in this fMRI study. This study was approved by the School of Psychology's Ethics Committee.

##### 2.1.2. Task

During the fMRI scan, participants were required to complete an associative visuospatial working memory task. The task comprised of blocks of memory trials and blocks of control trials (see Supplementary Fig. 4). Participants performed four runs of the task. Prior to entering the MRI scanner, participants were provided 12 practice trials of the memory task and 12 practice trials of the control task.

Within a memory trial, participants were presented with four abstract shapes presented at four horizontal locations across the screen. Three of the shapes were coloured, with participants needing to encode the shape, location and colour of the items (the encoding phase, 2500ms). The fourth shape was grey, with the shape and location of the grey item varying at random. The grey item was a placeholder for the location and did not need to be actively encoded. Participants were then presented with a scintillating, coloured mask (600ms) followed by a variable inter-stimulus interval (ISI). Each of the 6 trials within a block had a unique ISI ranging between 650 and 2150ms. This was followed by the recall phase, in which either a white shape cue or a white location cue was presented above four response options (the four possible colours, 2500ms). Participants had to recall the colour of the shape or at the location from the encoding phase. The recall phase was again followed by a coloured mask (600ms). Memory trials were separated by a variable inter-trial interval (ITI, varying between 900-2400ms). Memory trials spanned an average of 9s, with a total block duration of 54s. The trials timings and orders remained constant across all blocks.

Within a control trial participants were presented with three grey items and one coloured item above the four response options. Participants were simply required to respond to the colour of the item. This screen was presented for 2500ms followed by a 600ms mask. Control trials spanned an average of 4.5s including a variable ITI (650-2650ms), with a total block duration of 27s. Timings of the control trials were based on the timings for the encoding phase of the memory trials.

Within each of the four runs of the task, participants were presented with four blocks of six memory trials and four blocks of six control trials, presented in a counterbalanced order. Blocks were separated by a 30s rest period. Each run took a total of 468s (7 minutes, 48 seconds).

The order of the trials was pseudo-randomised across participants such that no participant encountered the same trial twice, and different participants were presented with different subsets of trials. Across all memory trials in a run (24) there were an equal number of shape and location cues and an even number of each response key. These were randomly separated across blocks but with the constraints that no more than four instances of a given cue (location, shape) and no more than three instances of a given response key could be presented in each block. Each memory block contained an equal number of shape and location cues, and an equal number of the four options as correct colour.

Supplementary Figure 4

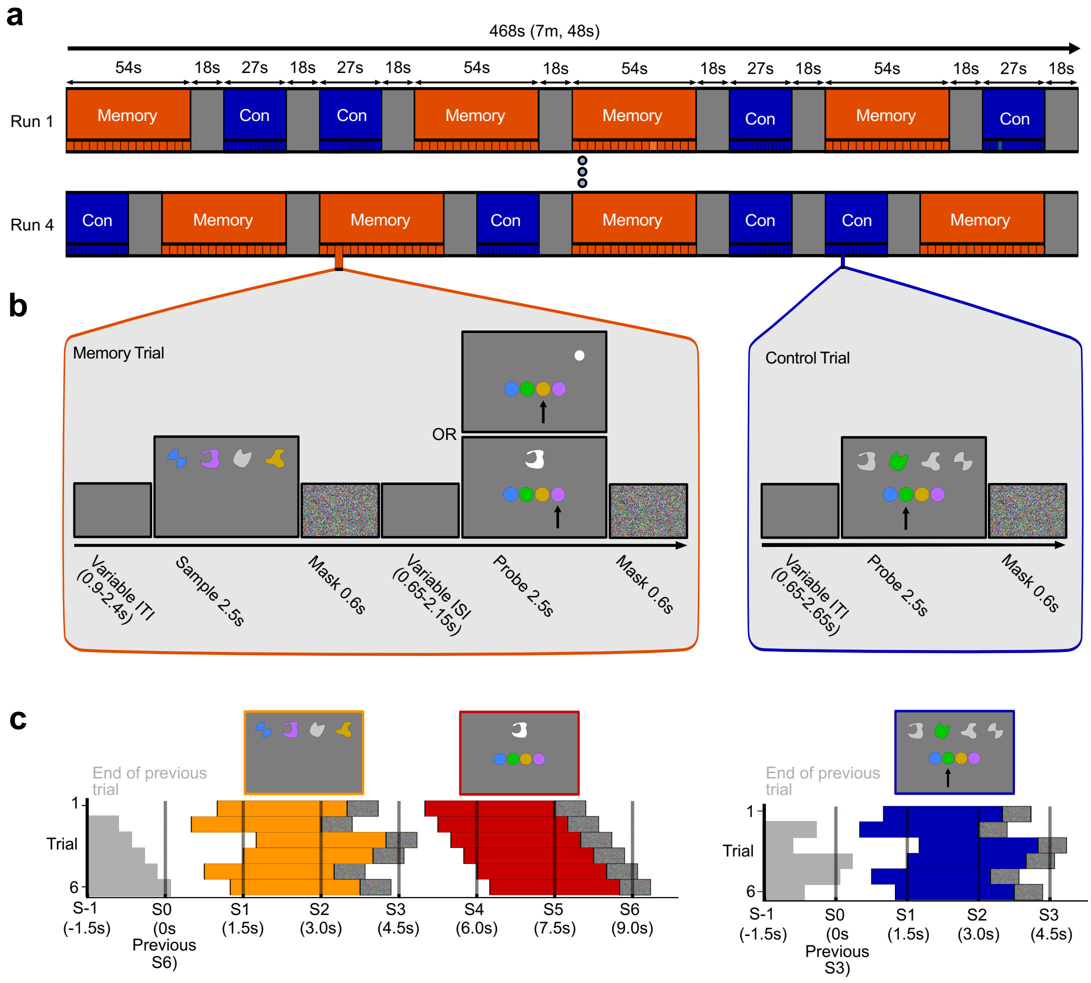


*Note:* (**a**) A depiction of two runs of the task, showing two of the counterbalancing options used in the study. (**b**) Depictions of a memory trial (left) and control trial (right). In the memory trial, participants may be presented with a location probe (top) or shape probe (bottom). The arrows show the correct answer and are for illustration purposes only. ITI = inter-trial interval, ISI = inter-stimulus interval. (**c**) Plots showing the timings of the stimuli in relation to sample acquisition for the memory trials (left) and control trials (right). Each row represents a trial within a block, starting with the top row. The textured bars represent the mask durations. S1, S2 etc refer to the trigger onsets within the trial. Grey lines show when then previous trial ended.

##### 2.1.3. fMRI data acquisition

Data were acquired on a 7T MR scanner (Phillips Achieva) using a 32 channel receive and 2 channel transmit head coil (Nova Medical).

An anatomical 3D T1-weighted PSIR acquisition was first acquired, TR = 6.3 s, voxel size = 0.7 mm isotropic, TE = 2.62 ms, flip angle = 5°, field of view = 157 x 235 x 235 mm. Functional (BOLD) images were acquired with a 2D gradient echo EPI sequence (multiband 2, SENSE $r = 2$). TR = 1500 ms, voxel size = 2 mm isotropic, TE = 25 ms, flip angle = 70°, field of view = 100 x 192 x 192 mm. Data was acquired over 50 axial slices; 96 × 96 voxels per slice. Functional images were acquired across all four runs of the task, with 314 volumes acquired within a run. Five dummy volumes were acquired and discarded prior to data collection.

##### 2.1.4. Data preprocessing and analysis.

Functional data were analysed with FSL [1]. Thermal noise was accounted for with NORDIC preprocessing [2] and top-up scans were applied to account for distortion. The data were motion corrected within scans using McFlirt motion correction, using linear interpolation. Drift correction was applied. The motion-corrected functional runs were aligned to the participants' anatomical scans using a full search and 12 degrees of freedom. Functional data were spatially smoothed (3D Gaussian, FWHM 5 mm), temporally high-pass filtered (cut-off 0.01 Hz) and FILM pre-whitening was applied.

Three levels of analysis were performed. The first analysed each individual run, with the second combining the four runs within a participant. The third level analysed the comparisons of interest across the group.

To find areas more active during the memory trials, we used the following contrast: “memory > control”. As this was a pilot localiser study with only five participants, with the aim to identify the region of MFC engaged in the memory task, we performed the comparison at the group level without any cluster or voxel-wise correction.

#### 2.2. Trial timings

During the fMRS study, the two timing solutions provided non-sequential orders of the 16 sampling points whilst ensuring that a unique ISI was present for each of the 16 memory trials and that the encoding and recall phases were sampled at different times within a trial. Four-hundred possible solutions were sampled, with the two chosen that had the highest combined sum of the square error (SSE) comparing the times of the sampling points across solutions, for both the encoding and recall phases (see Supplementary Eq. 1).

Supplementary Equation 1

$$\text{combined } SSE = \sum (E1 - E2)^2 + \sum (R1 - R2)^2$$

Where E1 and E2 are the vectors of sampling points for the encoding phases of solution 1 and 2 respectively, and R1 and R2 are the vectors of the sampling points for the recall phases of solution 1 and 2 respectively.

### 3. References


1. Jenkinson, M., Beckmann, C. F., Behrens, T. E. J., Woolrich, M. W. & Smith, S. M. FSL. *NeuroImage* **62**, 782–790 (2012).
2. Moeller, S. *et al.* NOise reduction with DIstribution Corrected (NORDIC) PCA in dMRI with complex-valued parameter-free locally low-rank processing. *NeuroImage* **226**, 117539 (2021).
